# Oral exposure to PET microplastics alters the pancreatic transcriptome – implications for the pathogenesis of type 1 diabetes

**DOI:** 10.1101/2024.11.05.622142

**Authors:** Karol Mierzejewski, Aleksandra Kurzyńska, Monika Golubska, Ismena Gałęcka, Jarosław Całka, Iwona Bogacka

## Abstract

Type 1 diabetes is a chronic autoimmune disease, the incidence of which has been steadily increasing in recent years, particularly among adolescents. The disease results from a combination of genetic and environmental factors that lead to the destruction of insulin-producing beta cells in the pancreas. Recently, the potential role of microplastics in the pathogenesis of various diseases has gained attention. Therefore, the aim of this study was to investigate the impact of PET microplastics on the pancreas using immature pigs as a model organism. The global transcriptomic profile of the pancreas was analyzed in piglets treated with either a low (0.1 g/day) or high dose (1 g/day) of PET microplastics for 4 weeks using RNA-Seq. The analysis revealed a dose-dependent effect of PET microplastics on gene expression. A low dose affected the expression of one gene, while a high dose impacted the expression of 86 genes. The differentially expressed genes, including immune cell markers, cytokines and chemokines, may activate the immune system in the pancreas in a way that is characteristic of the pathogenesis of type 1 diabetes. It suggests that oral exposure to PET microplastics may be a new risk factor for the development of this disease.

## Introduction

Diabetes mellitus (DM) is a long-term metabolic disease characterized by constantly elevated blood glucose levels, either due to insufficient insulin production or the body’s inability to effectively use the insulin it produces (1). This disease affects people of all ages, genders and regions, making it one of the leading causes of mortality and morbidity worldwide. Therefore, diabetes is considered one of the fastest growing health challenges of the twenty-first century and represents a significant financial burden for healthcare systems worldwide. An estimated 10.5% of the world’s population currently live with this disease (2).

Based on their pathogenesis, two main types of diabetes can be distinguished: type 1 diabetes (T1D) and type 2 diabetes (T2D) (3). The prevalence of T1D is estimated for approximately 10% of all diabetes cases worldwide and is often diagnosed in children and adolescents, although it can occur at any age. In contrast, T2D is much more common and accounts for around 90% of all diabetes cases, with an increasing incidence in adults as well as in children due to lifestyle factors such as obesity (4). Both types are characterized by a defect in the β-cells of the pancreas, which play a crucial role in insulin production. However, the causes of the dysfunction and death of the β-cells differ between the two types. In T1D, the insulin-producing β-cells are damage mainly by autoimmune mechanisms, leading to insulin deficiency (5). In T2D, the β-cells may be present but are unable to secrete sufficient insulin to compensate for insulin resistance in peripheral tissues such as muscle or fat tissue. Over time, β-cell function may continue to decline due to chronic insulin demand (6). Thus, the loss of functional β-cells is a crucial for the development of hyperglycemia and diabetes in both forms of the disease. The prevalence of diabetes is associated with certain risk factors, including lifestyle, diet, insufficient physical activity, obesity, alcohol consumption or genetic predisposition. In light of recent studies, it seems that exposure to microplastics may be another significant risk factor for diabetes development.

Microplastics (MPs) are small plastic particles, measured in size less than 5 mm. They are created by the chemical, physical or biological degradation of larger plastic parts (secondary microplastics) or are intentionally produced this small (primary microplastics) and are used, for example, as an additive in cosmetic products (7). Due to their ubiquity in water, food and air, microplastics have caused great concern about their impact on human health in recent years. Their presence has already been detected in human tissue, e.g. in the placenta, testicles, liver, brain or blood (8–10). Microplastic particles have been shown to influence numerous physiological processes, including the activation of the immune response, the triggering of inflammation and oxidative stress (11). There is also evidence that microplastics may cause metabolic diseases associated with insulin resistance by dysregulating the gut microbiota (12) or affecting blood glucose levels (13). In addition, our previous study has shown that PET microplastics alter the expression of miRNA in serum-derived extracellular vesicles (EVs), which are associated with insulin resistance, type II diabetes and the development of pancreatic cancer (14). Considering our previous findings and literature data, this study was conducted to determine the role of PET microplastics on the global transcriptomic profile of the pancreas using immature piglets as a model organism. The pig is considered as a valuable model for studying diabetes due to its physiological similarities to humans, particularly in pancreas structure, metabolic function, and pathophysiological responses (15). We have shown that PET microplastics affect the expression of genes involved in chemotaxis and immune cell activation in pancreases, which could be associated with the pathogenesis of type 1 diabetes.

## Materials and methods

### Animals

All experimental procedures were approved by the Local Ethics Committee of the University of Warmia and Mazury in Olsztyn (Decision No. 10/2020, dated February 26, 2020), and the study was conducted in compliance with the European Union Directive on the ethical treatment of animals used in experiments (EU Directive 2010/63/EU). The animals were housed in standard laboratory conditions, with free access to fresh water (ad libitum) and an age-appropriate diet. The temperature in the pens was maintained between 20-22°C, and humidity levels were kept between 55-60%. All plastic materials were removed from the animals’ surroundings, and stainless steel was used for the feeding and watering equipment. The farm equipment where the gilts were housed prior to the experiment was also stainless steel. Wood and bedding materials were used in the environment to ensure high welfare standards. The experiment lasted for four weeks and involved 15 eight-week-old gilts (Pietrain x Duroc) weighing approximately 20 kg. The animals were randomly assigned to three groups: 1) a control group (CTR, n = 5), which received empty gelatin capsules orally; 2) a low-dose group (LD, n = 5), which received a daily oral dose of 0.1 g/pig of PET microplastics (MPs) in gelatin capsules; and 3) a high-dose group (HD, n = 5), which received 1 g/pig of PET MPs per day in gelatin capsules. The capsules were administered one hour before the morning feeding. The plastic used in the experiment was semi-crystalline polyethylene terephthalate (PET) powder, supplied by Goodfellow Cambridge Ltd., UK (Cat. No ES306031/1). Details regarding the plastic particle have been described previously (16,17). The doses administered were selected based on the weekly human consumption of MPs reported in the literature (18,19) and adjusted to the weight of the gilts. After four weeks, the piglets were euthanized. The euthanasia protocol involved the administration of atropine (0.05 mg/kg i.m., Polfa, Poland), followed by xylazine (3 mg/kg i.m., Vet-Agro, Poland) and ketamine (6 mg/kg i.m., Vetoquinol Biowet, Poland). Approximately 20 minutes later, when the animals were fully anesthetized, an overdose of sodium pentobarbital (0.6 mL/kg i.v., Biowet, Poland) was administered. The cessation of vital functions was confirmed by the absence of a pupillary reflex, pulse, and respiration, after which pancreatic tissue samples were immediately collected for further analysis.

### RNA isolation, library preparation and sequencing procedure

Total RNA from 15 samples was extracted using the RNeasy Mini Kit (Qiagen, Germany) according to the manufacturer’s instructions. The purity and concentration of the isolated RNA was determined using a Tecan Infinite M200 plate reader (Tecan Group Ltd., Switzerland). RNA degradation was assessed using the Agilent Bioanalyzer 2100 (Agilent Technology, USA). The process of library preparation and sequencing has been described previously (20). In brief, libraries were prepared using the TruSeq Stranded mRNA LT Sample Prep Kit (Illumina, San Diego, CA, USA). The RNA was fragmented, and reverse transcribed into cDNA. The double-stranded cDNA fragments were labeled with specific adapters for each library. The resulting cDNA fragments were strand specific. Finally, the pooled libraries were sequenced on the Illumina NovaSeq 6000 platform with 2 × 150 bp paired-end (PE) sequencing.

### Quality control and genome mapping

The quality of raw paired-end reads was controlled using FastQC and Trimmomatic, and sequences were processed to (a) be at least 120 bp long, (b) have a PHRED scoreof > 20, and (c) be trimmed to the same length. High quality trimmed reads were aligned to the Sus_scrofa 11.1 genome assembly with reference to the ENSEMBL annotation (release 98) using the STAR Aligner (Spliced Transcripts Alignment to a Reference). The mapping results were indexed and sorted by coordinates. The gene expression values (read counts) were reconstructed by compiling the ballgown files and the prepDE.py script.

### Differentially expressed genes

The analysis of differentially regulated genes (DEGs) with a false discovery rate (FDR < 0.05) and fold change threshold (absolute log2FC > 1.5) was performed using DESeq2. Changes in gene expression patterns in the porcine pancreas after *in vivo* treatment with PET microplastics were determined by high-throughput transcriptome sequencing. The transcriptomic effects of PET microplastics treatment were investigated in three comparisons: low dose PET MPs (LD PET) versus control (CTR), high dose PET MPs (HD PET) versus CTR, and HD PET versus LD PET. Additionally, fragments per kilobase of transcript per million mapped reads (FPKM) were calculated as a normalized expression measure that depends on sequencing depth and genomic feature length. The enrichment of key biological processes and metabolic pathways in DEGs was identified using the enrichGO and enrichKEGG methods implemented in the ontology-based clusterProfiler R package. For functional enrichment analysis, the criteria were set as follows: Organism: pig; ontology categories: CC, MF, or BP; p-adjust value cut-off: 0.05; p-adjust method: BH. The R Bioconductor packages ggplot2, circlize, and GOplot were used to generate expression and functional profiles (21,22).

### Real-time PCR validation

Differentially expressed genes were randomly selected for validation by real-time PCR using the AriaMx real-time PCR system (Agilent Technology, USA) as previously described (23). Primer sequences for the reference and target genes (*CCL22, CXCL10, IL12B, TNFSF14)* were designed using Primer Express Software 3 (Applied Biosystems, USA). PCR reaction mixes with a final volume of 25 μl consisted of cDNA (4 ng), 300 μM of each primer, 12.5 μl of Power SYBR Green PCR Master Mix (Applied Biosystems, USA) and RNase-free water. The abundance of the tested mRNAs was calculated using the comparative Pfaffl method (24). The constitutively expressed *PPIA* and *RPLP* genes were implemented as reference genes, and the geometric mean values of the expression levels were used for analysis. Real-time PCR results were analyzed using Statistica software (version 13.1; Statsoft Inc. Tulsa, OK, USA) with Student’s t-test and expressed as means ± SEM. The results were considered statistically significant at p ≤ 0.05.

## Results

### The effect of PET microplastics on differential gene expression in pancreases

Treatment of piglets with a low dose of PET microplastics revealed only one differentially expressed gene (DEG) in the pancreas – *ACTG2*, which was downregulated (Fig. 1A). In contrast, treatment with a high dose of PET microplastics resulted in the identification of 86 DEGs, with 7 genes being downregulated and 79 upregulated (Fig. 1B, 2A). Gene Ontology (GO) annotation of biological processes (BP) included 533 terms, while 174 terms were classified under molecular functions (MF) and 94 under cellular components (CC). According to GO analysis, the DEGs involved in the pancreatic response to PET microplastics were associated with processes such as immune response (*CCR7, CCL17, TNFSF14, LAX1, IRF8, CCL22, CXCL9, CXCL10, LTB, C3, ENSSSCG00000056654, ENSSSCG00000036445)*, chemotaxis (*CCR7, CCL17, CCL22, CXCL9, CXCL10, ENSSSCG00000036445, DOCK2*), cytokine activity (*IL12B, CCL17, CCL22, LTB, CXCL9, CXCL10, ENSSSCG00000036445*), and B cell differentiation (*RAG1, CARD11, IL2RG, MS4A1, LYL1*) (Fig. 2B). KEGG pathway analysis indicated that the identified DEGs were involved in cytokine−cytokine receptor interaction (*IL12B, CCR7, CCL17, TNFSF14, TNFRSF18, CCL22, LTB, CXCL9, CXCL10, ENSSSCG00000036445, IL2RG*), Th1 and Th2 cell differentiation (*IL12B, RUNX3, CD3D, IL2RG, JAK3*), T cell receptor signaling pathway (*CARD11, PIK3CD, CD3D, LCP2*), Toll-like receptor signaling pathway (*IL12B, PIK3CD, CXCL9, CXCL10*), NF-kappa B signaling pathway (*TNFSF14, CARD11, LTB, TRAF1*), and the JAK-STAT signaling pathway (*IL12B, PIK3CD, IL2RG, JAK3*) (Fig. 2C). Detailed results of DEGs, GO, and KEGG analyses are provided in Tables S3 and S4 of the Supplemental Materials.

**Figure 1.**
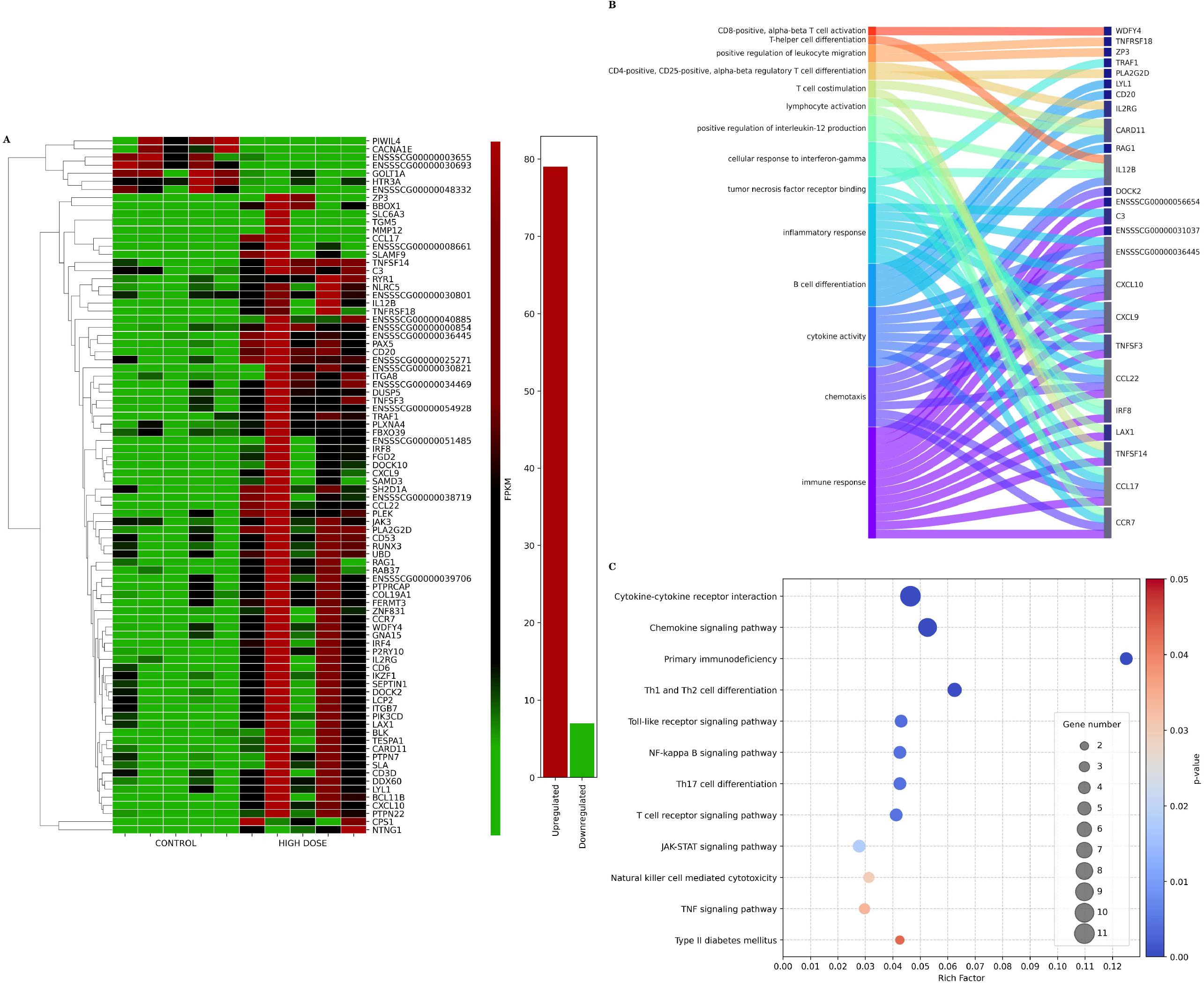
Volcano plots illustrating changes in gene expression profiles in porcine pancreas under the influence of PET microplastics. Plot **A** shows gene expression changes following a low dose of PET microplastics treatment, plot **B** shows changes after a high dose, and plot **C** compares gene expression between high and low doses of PET microplastics. The X-axis displays the logarithmic fold changes in expression (log2FC), while the Y-axis represents the - log10 transformed adjusted p-values (q-values). A horizontal dashed line marks the significance threshold for the adjusted p-value (0.05), and vertical dashed lines denote the fold change cut-off (absolute value of log2FC > 1.5). Each dot represents a gene’s expression level, with red dots indicating significantly overexpressed genes and green dots indicating significantly underexpressed genes (both with an adjusted p-value < 0.05). Grey dots represent genes with non-significant changes in expression.

**Figure 2.**
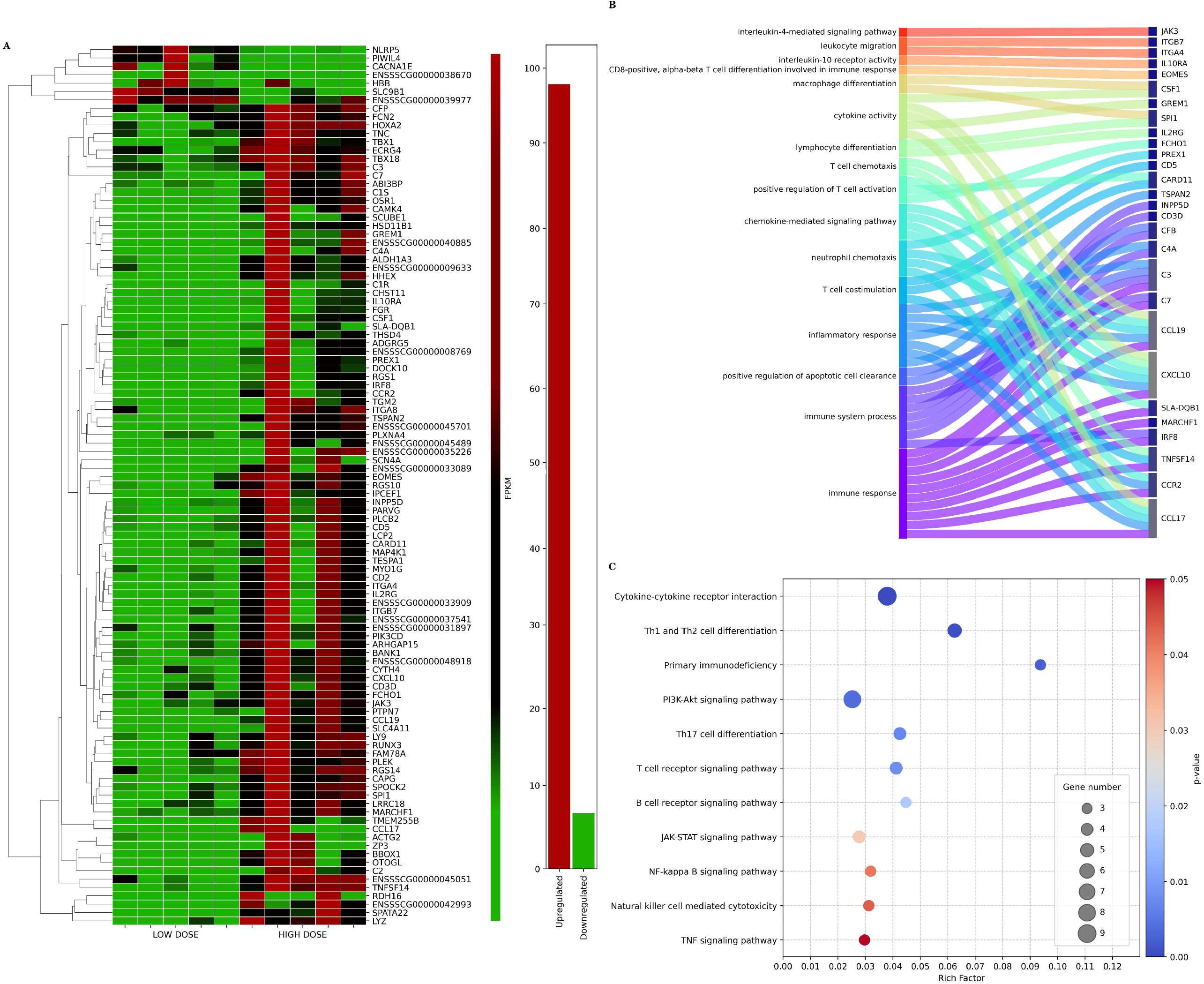
Gene expression analysis in porcine pancreas after exposure to high dose of PET microplastics. **A:** Heatmap showing the expression levels of differentially expressed genes in control and high-dose PET microplastics-treated groups. Rows represent individual genes, and columns represent samples from each treatment group (control vs. High dose). Colors indicate the level of gene expression, with red representing upregulated genes, green representing downregulated genes, and black indicating no significant change in expression. The dendrogram on the left clusters genes based on similarity in expression patterns. **B:** Sankey diagram illustrating enriched biological processes and pathways associated with the differentially expressed genes in the high dose of PET microplastics-treated group. This diagram shows how specific genes (on the right) are linked to various biological processes (on the left). Each pathway is represented by a different color, and lines connect genes to the biological processes in which they are involved. **C:** Bubble plot showing pathway enrichment analysis for the genes affected by high dose of PET microplastics. Each pathway is listed on the Y-axis, and the X-axis displays the “rich factor,” which represents the ratio of differentially expressed genes associated with each pathway to the total number of genes in that pathway. The color of each bubble indicates the statistical significance (p-value) of the enrichment, with a gradient from red (less significant) to blue (more significant). The size of the bubbles represents the number of genes associated with each pathway.

### Comparison between the low and high doses of PET MPs

This analysis identified 105 differentially expressed genes (DEGs) in the pancreases of gilts administered a high dose of PET microplastics (MPs), compared to those given a low dose. Of these, 7 genes were downregulated and 98 were upregulated (Fig. 1C, 3A). The Gene Ontology (GO) annotation of biological processes (BP) included 663 terms, while 195 terms were classified under molecular functions (MF) and 105 under cellular components (CC). Most of the processes were associated with immune system activation, including immune response (*CCL17, CCR2, TNFSF14, IRF8, MARCHF1, SLA-DQB1, CXCL10, CCL19, C7, C3*), follicular B cell differentiation (*IRF8, SPI1*), inflammatory response (*TSPAN2, CCL17, CCR2, C4A, CXCL10, CCL19, C3*), and T cell costimulation (*TNFSF14, CARD11, CD5*) (Fig. 3B). KEGG pathway analysis revealed that the DEGs were involved in the chemokine signaling pathway (*FGR, PLCB2, CCL17, CCR2, PIK3CD, CXCL10, CCL19, PREX1, JAK3*), cytokine−cytokine receptor interaction (*CCL17, CCR2, TNFSF14, CXCL10, IL10RA, CCL19, IL2RG, ENSSSCG00000009633, CSF*1), Th1 and Th2 cell differentiation (*RUNX3, SLA-DQB1, CD3D, IL2RG, JAK3*), and the PI3K−Akt signaling pathway (*PIK3CD, TNC, ITGA8, IL2RG, ITGA4, JAK3, CSF1, ITGB7)* (Fig. 3C). Detailed results of the DEGs, GO, and KEGG analyses are provided in Tables S3 and S4 of the Supplemental Materials.

**Figure 3.**
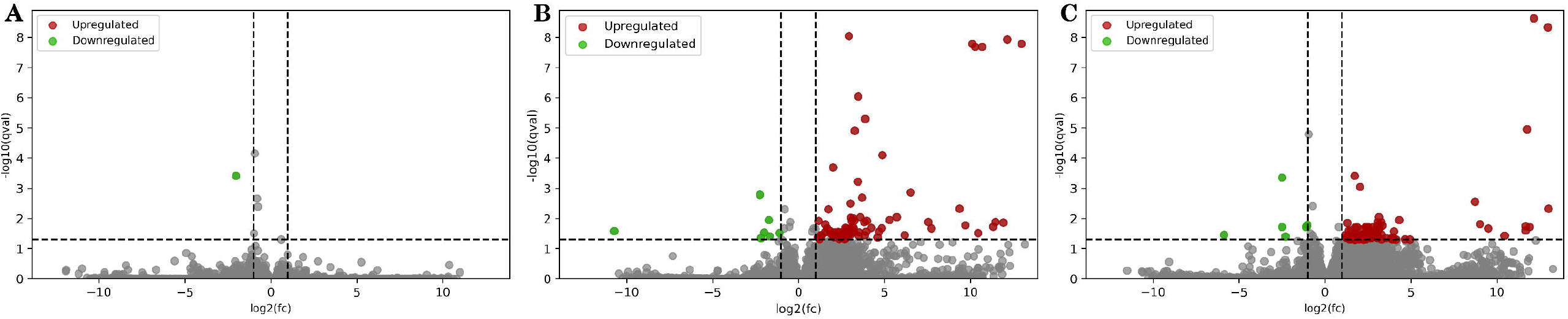
Gene expression analysis in porcine pancreas comparing the effects of high and low doses of PET microplastics. A: Heatmap displaying the expression levels of differentially expressed genes in the pancreas for both high dose and low dose PET microplastics treatment groups. Rows represent individual genes, and columns represent samples from each treatment group (low dose vs. high dose). Colors indicate the level of gene expression, with red representing upregulated genes, green indicating downregulated genes, and black showing genes with no significant change in expression. The dendrogram on the left clusters genes based on similarity in their expression patterns across both doses. **B**: Sankey diagram illustrating enriched biological processes and pathways linked to the differentially expressed genes when comparing high and low doses of PET microplastics. This diagram shows how specific genes (on the right) connect to biological processes (on the left), such. Each pathway is represented by a different color, with lines connecting genes to the biological processes they influence, highlighting differences in pathway involvement between the two doses. **C:** Bubble plot representing pathway enrichment analysis for genes with differential expression between high dose and low dose PET microplastics exposure. Pathways are listed on the Y-axis, while the X-axis shows the “rich factor,” which reflects the ratio of differentially expressed genes in each pathway to the total number of genes in that pathway. The color gradient of each bubble indicates the statistical significance (p-value) of the enrichment, with a range from red (less significant) to blue (more significant). The size of the bubbles represents the number of genes associated with each pathway.

### Real-time PCR

The analysis revealed that CCL22, CXCL10, IL12B, TNFSF14 mRNA abundance were upregulated in the pancreas after treatment of the piglets with a high dose of PET microplastics. Real-time PCR expression patterns of the tested DEGs agreed with RNA-Seq results (Supplemental Fig. 1).

## Discussion

This study provides evidence for a new potential risk factor, PET microplastics, which can significantly affect physiological processes in the pancreas. Considering that the average human plastic intake is estimated to be between 0.1 g and 5 g per week (18), two doses of PET microplastics – 0.1 g/day and 1 g/day – were considered in our *in vivo* experiment. The lower dose of PET microplastics altered the expression of one gene, whereas the higher dose altered the expression of 86 genes. The fact that the low dose had a minimal effect on the transcriptomic profile of the pancreas, while the high dose had a significant effect, suggests a dose-dependent response at the transcriptomic level. A comparison between the two doses revealed 105 differentially expressed genes, most of which were involved in the regulation of immune responses and immune cell differentiation. This indicates that the pancreas may tolerate or cope with low exposure to PET microplastics without significant changes in gene expression. However, at higher concentrations, the organism’s ability to cope is overwhelmed, leading to substantial transcriptomic changes. This may reflect a threshold beyond which microplastic exposure triggers stress response, inflammation, or cell dysfunction, resulting in toxic effects on the pancreas. Consequently, controlling the amount of microplastics ingested could be crucial in mitigating their harmfulness.

Our study shows that PET microplastics strongly affect the immune response in the pancreas. KEGG analysis revealed that the differentially regulated genes are involved in the regulation of cytokine and chemokine activity, lymphocyte differentiation, and TLR4 and NF-kappa B signaling pathways. It is well known that diabetes, like most autoimmune diseases, is characterized by an overproduction of pro-inflammatory cytokines. Our findings show that PET microplastics increase the expression of *IL-12*β in the pancreas. IL-12 has been reported to play a crucial role in the development and pathogenesis of autoimmune diseases by driving the recruitment of inflammatory cells (25). In non-obese diabetic (NOD) mice, increased expression of this cytokine was found in islet cells, in parallel with the destruction of β-cells (26). Another study in NOD mice indicated that IL-12 plays a significant role in the transition from non-destructive to destructive insulitis (27,28). In addition to IL-12β, our study shows that PET microplastics upregulated the expression of various members of the TNF superfamily, such as *TNFSF3* (LTβ), *TNFSF14* (LIGHT) and *TNFRSF18* (GITRL). These pro-inflammatory cytokines affect both the immune system and β-cell function by binding to their corresponding receptors (TNFRSF) (29). TNFSF3 receptors, which are predominantly expressed on immune cells and lymphoid stromal cells, contribute to the ectopic formation of tertiary lymphoid organs (TLOs) that trigger chronic inflammation and autoimmune responses (30). These processes facilitate persistent immune activation and β-cell destruction, promoting the development and progression of type 1 diabetes (31,32). Additionally, TNFSF14 has been linked to immune-mediated destruction of β-cells in diabetic mice (33), and elevated levels of this cytokine have been observed in diabetic patients (34). TNFRSF18, in turn, exacerbates autoimmune diabetes by selectively activating aggressive T cells while suppressing Tregs. It has been shown that blocking TNFRSF18 expression can prevent the onset of diabetes (35). These findings suggest that PET microplastics may increase the risk of β-cell damage and contribute to the development of type 1 diabetes by upregulating key diabetogenic cytokines such as *IL-12*β, *TNFSF3, TNFSF14* and *TNFRSF18*.

This study shows that a high dose of PET microplastics upregulate the expression of several chemokines in the pancreas, including *CXCL10, CXCL9, CCL17*, and *CCL22*. Chemokines are specific cytokines that mediate the chemotaxis of immune cells by binding to surface receptors. Elevated levels of CXCL10 have been found in the serum of patients with newly diagnosed type 1 or type 2 diabetes and in patients at high risk of developing the disease (36–39). Furthermore, studies in mouse models have shown that CXCL10 is overexpressed in the pancreatic islets even before insulitis is detectable (40,41). CXCL9, in conjunction with CXCL10, also influences the mass and function of β-cells, and deletion of CXCR3 in mice delays the onset of type 1 diabetes (42). Thus, PET microplastics may activate immune responses in the pancreas and increase the risk of type 1 diabetes. Interestingly, we also observed increased expression of *CCL17* and *CCL22*, which act via the CCR4 receptor, predominantly expressed on Tregs. These chemokines are known to protect pancreatic β-cells from autoimmune attack in type 1 diabetes. CCL17 and CCL22 recruit Tregs to the islets, reducing autoreactive CD8+ T cells and mitigating β-cell destruction (43). The simultaneous upregulation of chemokines that promote (*CXCL10, CXCL9*) and protect against (*CCL17, CCL22*) type 1 diabetes suggests a complex immune response to PET microplastics. Initially, microplastics may promote β-cell destruction, but tissue injury could trigger a feedback mechanism that produces protective chemokines, such as CCL17 and CCL22, to limit further damage.

The current research indicates that PET microplastics upregulate the expression of several markers associated with immune cells in pancreatic tissue, such as *CD3D* and *CD6*, indicating T cell infiltration. The increased expression of *RUNX3* and *LCP2*, which support cytotoxic CD8+ T cell differentiation and regulate TCR signaling, further supports the notion that PET microplastics promote T cell infiltration (44,45). Furthermore, PET microplastics upregulated the expression of *JAK3*, a key enzyme in the JAK-STAT signaling pathway involved in regulating immune cell activity. JAK3, in combination with the IL-2 receptor, which was also upregulated in this study, plays a critical role in T cell development and the progression of type 1 diabetes (51). Inhibition of JAK3 has been shown to protect against diabetes onset in mouse models, highlighting its importance in diabetes pathogenesis (52).

Though T cells are largely responsible for β-cell destruction in type 1 diabetes, there is growing evidence that B cells also play a role in the pathogenesis of the disease. Research has shown that depleting B cells delays the progression of type 1 diabetes in newly diagnosed patients (46). In our study, PET microplastics increased the expression of *CD20* (MS4A1) as well as other markers important for B cell activation, such as *PAX5* and *BLK* (47). The presence of these markers suggests that PET microplastics might contribute to pathological processes such as autoimmune pancreatitis or type 1 diabetes.

Our study shows an increased expression of interferon regulatory factors (IRFs), particularly *IRF4* and *IRF8*. It is worth noting that both factors play a crucial role in the pathogenesis of type 1 diabetes by modulating interactions between β-cells and immune cells (32). Overactivation of IRF4 has been associated with increased secretion of diabetogenic cytokines by CD4+ T cells, promoting the development of type 1 diabetes (48,49). IRF8, in turn, is essential for the maturation of dendritic cells (DCs) and the production of type I interferons, which may exacerbate autoimmune attacks on β-cells (32,50).

All the above evidence suggest that PET microplastics trigger immune response genes in a manner characteristic of type 1 diabetes. Additionally, our previous study demonstrated that PET microplastics disrupt the pancreatic metabolome and may contribute to the dysfunction and potential damage of pancreatic β-cells through mechanisms involving oxidative stress, inflammation, and apoptosis (53). Moreover, we observed an increase in blood insulin levels, while blood glucose levels remained within the normal range, which is a known compensatory response of β-cells during the initial stages of dysfunction and could precede the onset of full-blown diabetes (54). Together, these studies suggest that PET microplastics affect both the metabolome and transcriptome of the pancreas, potentially impairing β-cell function which may accelerate the development of metabolic disorders such as diabetes.

In conclusion, our study suggests that orally ingested PET microplastics affect the transcriptomic profile of the pancreas in a dose-dependent manner. The upregulation of JAK3, cytokines, chemokines, and immune-related genes points to an inflammation- and immune-driven mechanism that may contribute to β-cell damage and possibly the development of type 1 diabetes. These findings offer a comprehensive insight into the biological processes affected by PET microplastics and provide a basis for further investigations into their role in the pathogenesis of type 1 diabetes.

## Acknowledgement

We would like to thank MSc Kamil Markuszewski for his help in preparing the visualizations of RNA-Seq results.

## Author contributions

Conceptualization: K.M., I.B.; Data curation: K.M.; Formal analysis: K.M.; Funding acquisition: K.M.; Investigation: K.M., A.K., M.G. I.G., J.C.; Methodology: K.M., M.G., I.G. J.C.; Project administration: K.M.; Resources: K.M., A.K., M.G.; Software: K.M.; Supervision: I.B.; Validation: M.G.;Visualization: K.M.; Writing: original draft: K.M.; Writing: review and editing: K.M., I.B., All authors have read and agreed to the published version of the manuscript.

## Data availability statement

The raw data generated for this study can be found in the European Nucleotide Archive (ENA) with the accession number PRJEB80780.

