## Supplementary material for "Oral exposure to PET microplastics alters the pancreatic transcriptome – implications for the pathogenesis of type 1 diabetes": Supplemantal Materials: Supplemental data description.docx

**Supplemental Fig. S 1**

Real-time PCR validation of RNA-seq results for random DEGs. Validation was performed for *CCL8*, *IL-1β*, *CCL4*, *OXTR* and *CSF3*.

**Supplemental Table S 1**

Genes identified in the porcine pancreas after treatment with (A) low dose of PET microplastics, (B) high dose, and (C) comparison between high and low dose of PET microplastics.

**Supplemental Table S 2**

Results of Gene Ontology enrichment analysis of DEGs significantly modulated in the porcine pancreas after treatment with (A) low dose of PET microplastics, (B) high dose, and (C) comparison between high and low dose of PET microplastics.

**Supplemental Table S 3**

Results of KEEG enrichment analysis of DEGs significantly modulated in the porcine pancreas after treatment with (A) low dose of PET microplastics, (B) high dose, and (C) comparison between high and low dose of PET microplastics.
