## Supplementary figures and images for "Oral exposure to PET microplastics alters the pancreatic transcriptome – implications for the pathogenesis of type 1 diabetes"

### Supplemental Figure S1.tif

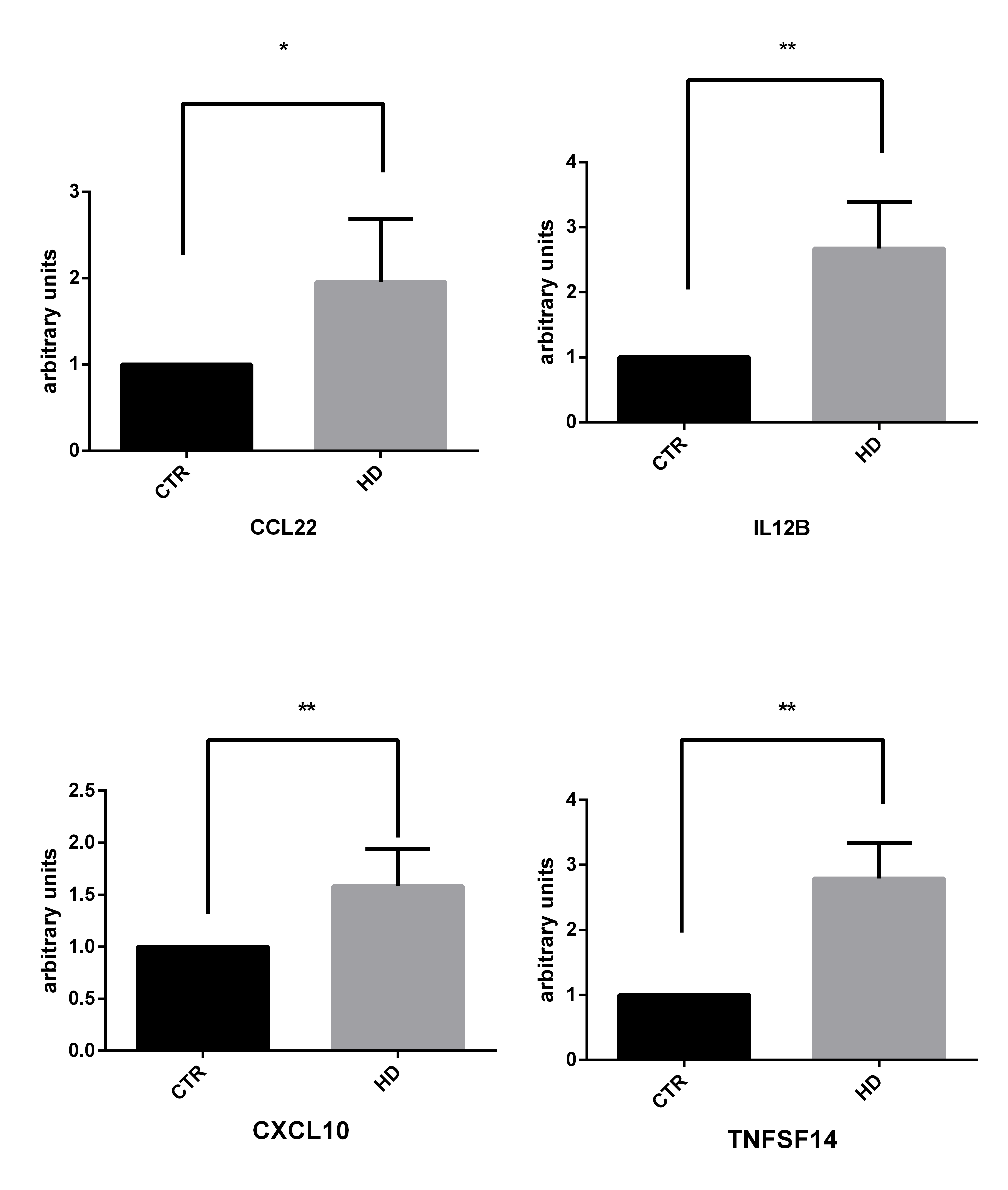
